# Intersegmental coordination of the central pattern generator via interleaved electrical and chemical synapses in zebrafish spinal cord

**DOI:** 10.1101/2021.09.07.459336

**Authors:** Lae Un Kim, Hermann Riecke

## Abstract

A significant component of the repetitive dynamics during locomotion in vertebrates is generated within the spinal cord. The legged locomotion of mammals is most likely controled by a hierarchical, multi-layer spinal network structure, while the axial circuitry generating the undulatory swimming motion of animals like lamprey is thought to have only a single layer in each segment. Recent experiments have suggested a hybrid network structure in zebrafish larvae in which two types of excitatory interneurons (V2a-I and V2a-II) both make first-order connections to the brain and last-order connections to the motor pool. These neurons are connected by electrical and chemical synapses across segments. Through computational modeling and an asymptotic perturbation approach we show that this interleaved interaction between the two neuron populations allows the spinal network to quickly establish the correct activation sequence of the segments when starting from random initial conditions and to reduce the dependence of the intersegmental phase difference (ISPD) of the oscillations on the swimming frequency. The latter reduces the frequency dependence of the waveform of the swimming motion. In the model the reduced frequency dependence is largely due to the different impact of chemical and electrical synapses on the ISPD and to the significant spike-frequency adaptation that has been observed experimentally in V2a-II neurons, but not in V2a-I neurons. Our model makes experimentally testable predictions and points to a benefit of the hybrid structure for undulatory locomotion that may not be relevant for legged locomotion.

## 1 Introduction

Animal locomotion requires the repetitive activation of muscles over a range of speeds that is adaptively varied to achieve a variety of goals including steady motion, turns to avoid obstacles, sudden escapes, and the like. Substantial portions of the motor programs performing these motions are generated within the spinal cord itself, with central pattern generators playing a key role.

In animals like lamprey, leech, and fish larvae the basic motion consists of an undulation wave traveling along the body. A striking experimental observation in the undulatory motion of various species is that the timing of the activity of the different segments of the animal depends only weakly on the frequency of the motion (Williams, 1992). More precisely, the delay in the oscillatory motion from one segment to the next is not given by a fixed time delay determined by synaptic time constants or axonal propagation speed but rather by a fraction of the period that is quite independent of the frequency of the wave. Importantly, this enables the mode of the physical motion of the animal’s body, i.e. the undulation wavelength, to stay almost the same across a range of frequencies. A fundamental question that arises therefore in the study of the spinal cord locomotor control is how the circuit is able to limit the dependence of the intersegmental phase difference (ISPD), i.e. the phase shift between successive segments, on the swimming frequency.

Fish and fish larvae often swim in spurts. The initiation of such a spurt requires that the spinal cord can quickly establish the correct phase relationships between the different segments, independent of their previous activity. If the organization of the phases were to take multiple oscillation periods, the swimming motion would initially be quite disorganized and ineffective.

The steady-state phase relationship of spinal-cord models has been studied extensively and comprehensively in the context of coupled-oscillator theory, where each segment corresponds to an individual oscillator that is coupled to its nearest and possibly further neighbors (Kopell and Ermentrout, 1986; Kopell et al., 1991; Cohen et al., 1992). For the commonly considered case of weakly interacting oscillators the coupling arises through the corresponding phase differences. For bidirectionally coupled oscillators it has been shown that the phase difference can be frequency-independent if the coupling function does not depend on the oscillation period (Kopell and Ermentrout, 1986; Kopell et al., 1991; Cohen et al., 1992). This has been obtained explicitly in some models that were formulated in terms of firing rates (Williams, 1992; Varkonyi et al., 2008). In general, however, the frequency-independence turns out not to be robust (Varkonyi et al., 2008).

The duration of the transient that is required to establish steady swimming seems not to have been addressed much so far. Within the weak-coupling framework, in which - due to the averaging over a period - the coupling between segments depends only on the phase differences (Kopell and Ermentrout, 1986), the ordering of the phases occurs formally on a time scale that is much longer than the period of the individual oscillations. This framework can therefore not reliably be used to investigate how coherent swim motion is established within a few oscillation cycles and one needs to use an approach that does not rely on this averaging procedure.

Classically, two classes of models have been proposed for the organization of spinal cord circuits (McLean and Dougherty, 2015). Particularly for legged locomotion, e.g. in mammals, a multilayer hierarchical structure has been proposed in which the timing of the motion is controlled by neurons that exclusively make higher-order connections to other interneurons and neurons in the brain stem, while the last-order projections to the relevant motor pools are exclusively made by neurons in a separate layer (McCrea and Rybak, 2008). This structure is especially motivated by the observation of so-called non-resetting perturbations in which the suppression of last-order neurons leads to a brief failure in the activation of the motor neurons but has no impact on the timing of subsequent activations, since it is exclusively controlled by the higher-order neurons. For simpler, axial locomotor circuits, as they arise, e.g., in lamprey and zebrafish larvae, a single-layer structure has been proposed in which the last-order neurons driving the motor pool also make higher-order connections to other interneurons, which control the timing of the activations (Kozlov et al., 2009).

Based on recent anatomical (Menelaou et al., 2014) and physiological studies (Menelaou and McLean, 2019) in zebrafish larvae a new, hybrid model has been proposed in which the excitatory last-order neurons not only drive the motor pool, but also project to other excitatory interneurons, as would be the case in a single-layer architecture. As in a multi-layer model, however, these neurons are comprised of two populations, termed V2a-I and V2a-II, which have analogs in the spinal cord of mouse (Hayashi et al., 2018). They differ in their anatomical and physiological properties, particularly in the range of their projections (Menelaou et al., 2014), the relative strengths of their last-order vs higher-order connections, and their mode of spiking (Menelaou and McLean, 2019).

Moreover, it was shown that the V2a neurons are connected via chemical synapses as well as electrical gap junctions (Menelaou and McLean, 2019). Interestingly, the gap junctions, which connect neurons of the same type (V2a-I) in different segments, appear to operate highly asymmetrically, with neurons essentially affecting only neurons that are located more caudally. Asymmetric coupling by gap junctions has been observed in other systems, albeit to a smaller degree (Nolan et al., 1999; Apostolides and Trussell, 2013; Sevetson and Haas, 2015). Thus, the coupling coefficient, which characterizes the voltage change induced in one V2a-I neuron by a voltage change in another V2a-I neuron, is different when going from neuron 1 to neuron 2 than vice versa. In contrast, the connections between V2a-I and V2a-II neurons in different segments are dominated by chemical synapses. So far, the function of these two populations of neurons and the role of the different types of connections are only poorly understood.

Here we investigate the role of the two neuron populations and the two types of connections in supporting swimming motion, with particular emphasis on the possible role of the connections *between* the two populations, which are characteristic of the hybrid model. We focus, in particular, on two aspects: on the ability of the system to quickly initiate the appropriate wave motion when swimming activity is initiated from random initial conditions and on the dependence of the inter-segmental phase differences on the swimming frequency. We find that in chains of V2a-I neurons that are only connected via gap junctions the requirement of fast initiation of waves that propagate reliably can only be satisfied for unphysiologically strong gap-junction coupling. Including, however, the appropriately timed additional excitatory input from V2a-II neurons *via* excitatory chemical synapses reduces the required gap-junction coupling to realistic values. Moreover, the spike-frequency adaptation of the V2a-II neurons provides a simple mechanism to limit the change of the intersegmental phase difference with the frequency of swimming.

## 2 Waves in Gap-Junction Coupled Chains of V2a-I Neurons

We are interested in the wave-like activation of successive segments of a long spinal cord. Motivated by the work of Menelaou and McLean (2019) we focus on the two types of excitatory V2a neurons. The V2a-I neurons in different segments are coupled among each other predominantly by gap junctions, while the V2a-II neurons receive input from upstream V2a-I neurons predominantly via glutamatergic chemical synapses and also excite their downstream V2a-I neighbors via such synapses (Figure 1). We will refer to the most rostral V2a-I neuron as the leading neuron and the rest as follower neurons. Note that in our model the V2a-II neuron in the leading segment receives glutamatergic input from the V2a-I neuron in the same segment. Here we will ignore the lateralization of the spinal cord and omit the inhibition between neurons on its left and right side (Kozlov et al., 2009).

**Figure 1:**
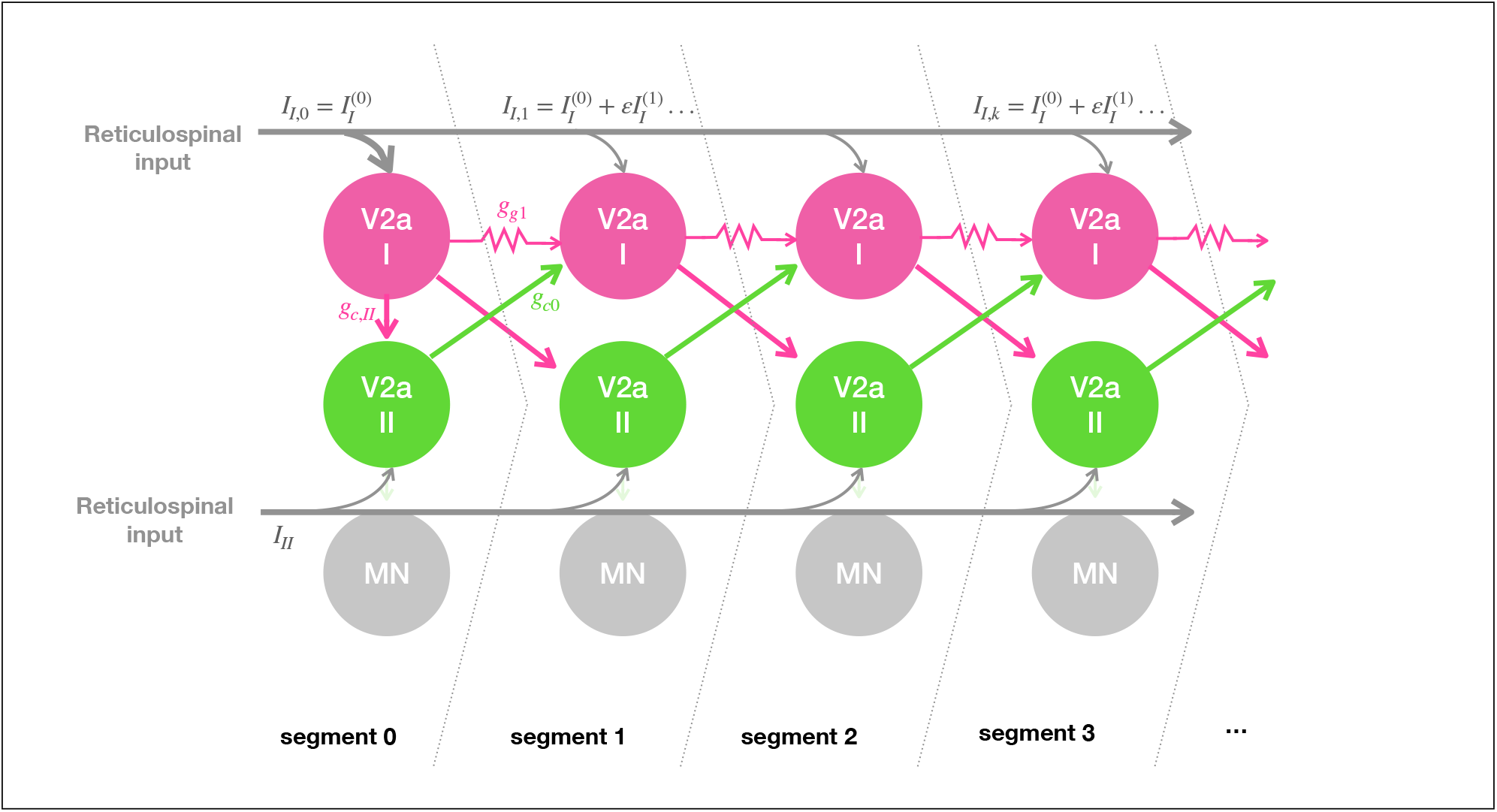
Network structure of the model. The V2a-I and V2a-II neurons form an interleaved network. Both types of neurons receive reticulospinal input. V2a-I neurons are connected unidirectionally via gap junctions. Connections between V2a-I and V2a-II neurons are via glutamergic chemical synapses. Note that the V2a-II neuron in the 0^*th*^ segment receives its V2a-I input from the *same* segment, which is an exception to the interleaved connection pattern.

Both types of neurons receive input from the reticular formation in the brain stem (Gahtan and O’Malley, 2003). These brainstem neurons have varying projection lengths, such that the shortest ones only reach the first couple of segments, while the longest ones reach up to two thirds of the whole spinal cord (Thiele et al., 2014). Thus, the total amount of input that the V2a neurons receive is likely not uniform along the spinal cord and depends on their location in the spinal cord. For the animal to go into an organized swimming motion quickly it is important that it reliably establish a stable phase-locked state within very few oscillations (Fig.2). Moreover, for the swimming motion to be consistent across different swimming speeds the intersegmental phase difference (ISPD) should not vary substantially with frequency.

**Figure 2:**
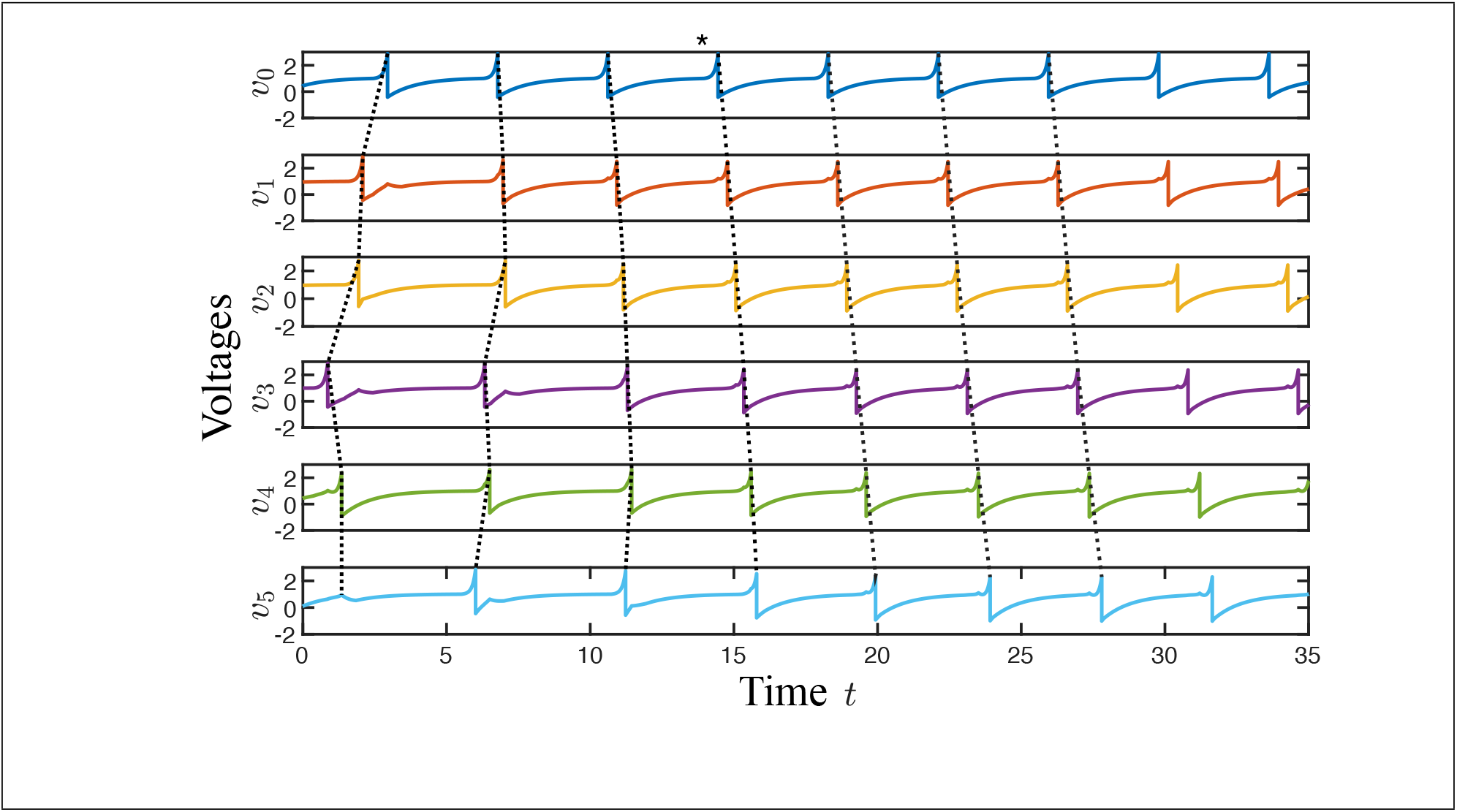
Wave-like excitation of a long spinal cord. A chain of length *N* = 6 of V2a-I neurons is simulated with random initial conditions. The correct ordering of the action potentials is achieved by the fourth cycle around *t* = 14 (marked with a star). The black dashed line connects corresponding spikes as a guide to the eye.

To gain insight into the mechanism allowing these dynamics and to obtain guidance for numerical simulations we choose very simple models for the neurons, which allow an analytical perturbation approach. At the same time, the prominent presence of gap-junction coupling requires that the voltage evolution during the action potential is approximated reasonably well. We therefore use the piecewise defined neuron model put forward by Chow and Kopell (2000), here written in dimensionless form,

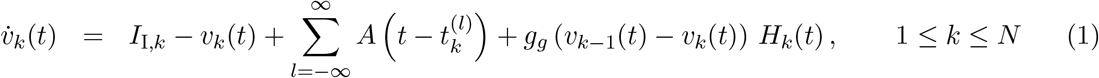

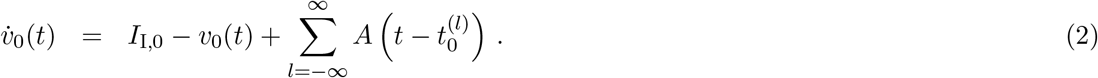

Here the action potentials are produced by the explicitly time-dependent exponential spiking current *A_S_*(*t*) and the recovery current *A_R_*(*t*),

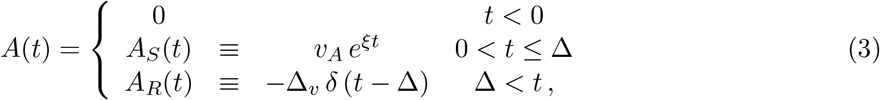

which are triggered whenever the scaled voltage *v*(*t*) reaches the threshold *v_T_* = 1 from below. In (1) this occurs for neuron *k* at the times 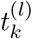. The spike width in this model is given by Δ and the recovery current that terminates the spike is modeled with a Dirac *δ*-function, which leads to a relative reset of the voltage by Δ_*v*_,

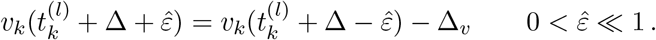

Menelaou and McLean (2019) report that the gap junctions are essentially unidirectional, such that the *k^th^* V2a neuron receives input from the (*k* – 1)^*th*^ neuron, but not vice versa. This may reflect that the gap junctions are located on the axons projecting caudally from each neuron; an action potential traveling along the axon can then drive current into the soma of the coupled neuron and excite there an action potential, but a somatic action potential in that neuron might not trigger a back-propagating action potential on the axon for lack of a spike-initiation zone on the axon near the gap junction. This suggests that the gap junction is effectively only carrying significant current during the axonal action potential, during which the axial voltage gradients are large. In-between action potentials the axon is essentially passive and does not couple the cells strongly. In our simulations we include this aspect by turning the gap junction on only during the spike in the upstream neuron and for a duration Δ thereafter, which is indicated by the function

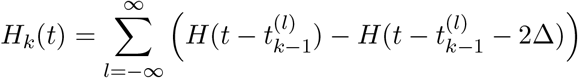

in (1). Here *H*(*t*) is the Heaviside step function. The simulations show that outside that interval the gap-junction current is very small (cf. Fig. 3 below). In our analytical treatment in Section 2.1 we omit this detail and have the gap-junction current active at all times (*H_k_* = 1).

**Figure 3:**
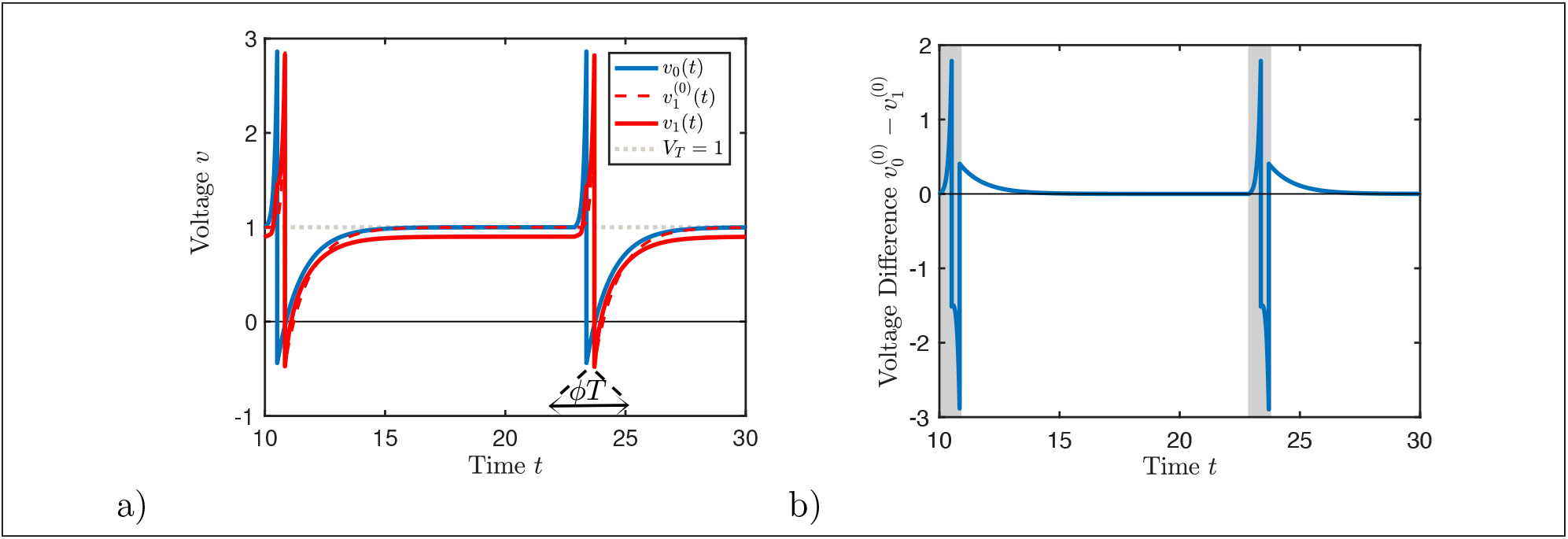
Voltages *v_k_*(*t*) of a chain of electrically coupled V2a-I neurons. (a) The voltage *v*_0_(*t*) of the leader neuron (blue) is given by the 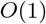-solution *V*^(0)^(*t*), while the voltage *v*_1_(*t*) of the follower neuron (red) has an 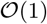-contribution 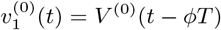 as well as an 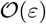-contribution 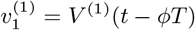, both delayed by *ϕT* relative to the leader neuron. The threshold *v_T_* for the onset of the spiking current is indicated by the grey dashed line. (b) The normalized gap junction current is given to 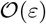 by *g_g_* (*V*^(0)^(*t* + *ϕT*) – *V*^(0)^(*t*)). In the simulations it is active only during the shaded area (cf. *Θ_k_*(*t*) in (1)). Outside that time interval the gap-junction current is very small. *I*_*I*,0_ is set for *f* = 77.7 (*T* = 12.9) and Δ*I* = 0.04.

### 2.1 Asymptotic Analysis for Weak Coupling

We first consider an infinite chain of V2a-I neurons that are coupled by gap junctions only. We are interested in a periodic traveling wave with frequency *f* ≡ 1,000/*T*. While all quantities are actually dimensionless, we have chosen parameters such that time can be thought of as measured in ms and frequency in Hz. In such a wave the voltage in segment *k* is given by

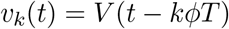

**Table 1:**
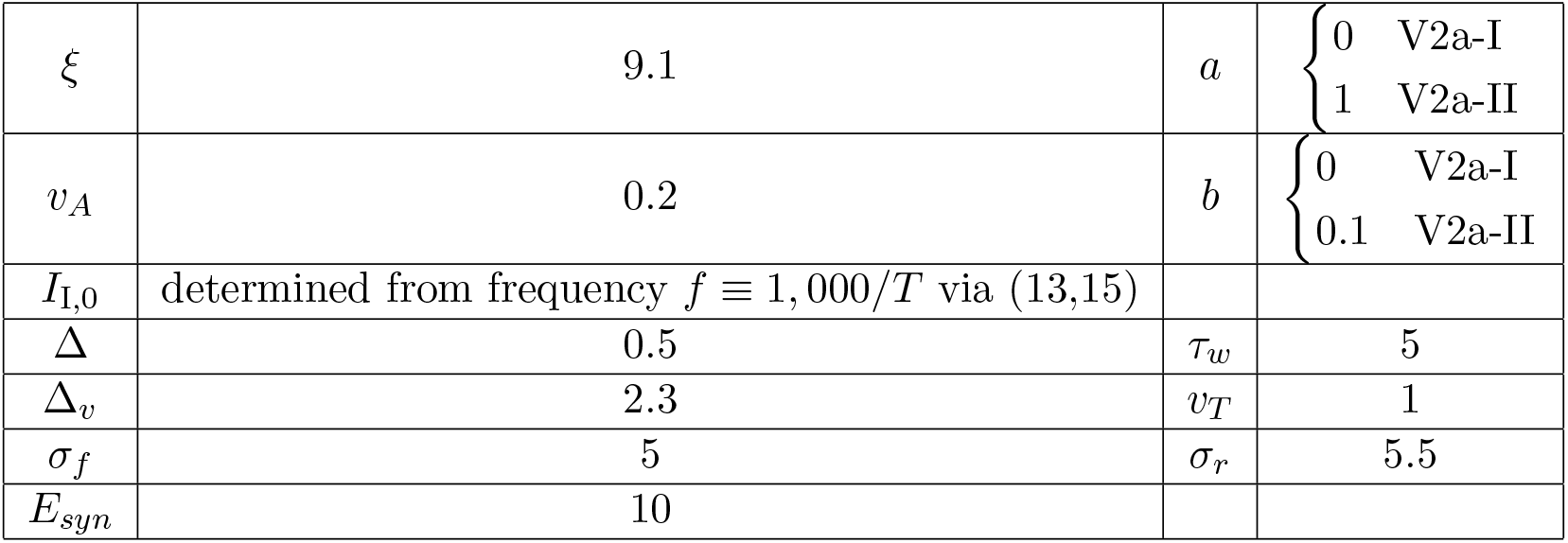
Parameter values for V2a-I neurons (cf. (1,3)) and for V2a-II neurons (cf. (21,22)) throughout the paper, unless otherwise specified.

|  |  |  |  |
| --- | --- | --- | --- |
| $\xi$ | 9.1 | $a$ | $\begin{cases} 0 & \text{V2a-I} \\ 1 & \text{V2a-II} \end{cases}$ |
| $v_A$ | 0.2 | $b$ | $\begin{cases} 0 & \text{V2a-I} \\ 0.1 & \text{V2a-II} \end{cases}$ |
| $I_{I,0}$ | determined from frequency $f \equiv 1,000/T$ via (13,15) | | |
| $\Delta$ | 0.5 | $\tau_w$ | 5 |
| $\Delta_v$ | 2.3 | $v_T$ | 1 |
| $\sigma_f$ | 5 | $\sigma_r$ | 5.5 |
| $E_{syn}$ | 10 | | |

with *ϕ* being the phase shift between adjacent segments (ISPD). For simplicity we take the origin in time such that the end of the *l^th^* spike in segment *k* occurs at time *t* – *kϕT* = *lT, l* integer, and consider *V*(*t*) only for 0 ≤ *t* ≤ *T*. Due to the piecewise definition of the time dependence of *A*(*t*) and the discontinuity generated by its recovery current (cf. (3)) the periodic solution *V*(*t*) satisfies

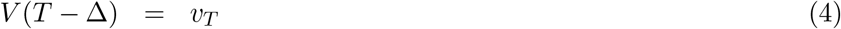

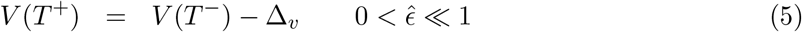

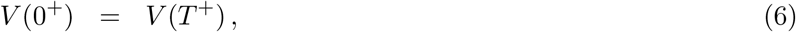

with 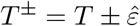 for 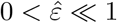.

To make analytic progress we assume the gap-junction coupling to be weak,

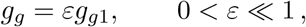

and expand the membrane voltages *V*(*t*) of the V2a-I neurons also in *ε*,

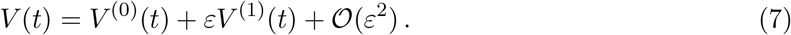

Because of the threshold and reset conditions (4,5) it is preferable to keep the period *T* fixed and expand the reticular input currents *I_I_* that are needed to obtain that period,

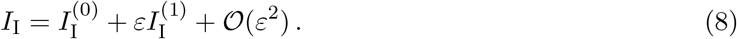

At 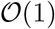 this leads to

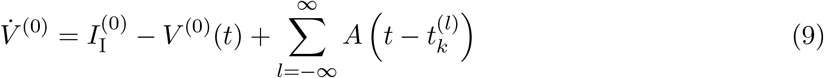

with

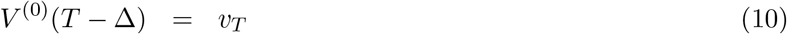

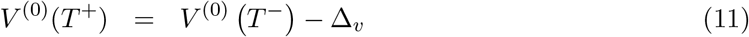

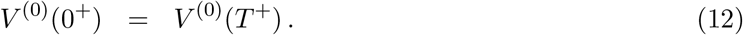

For given current 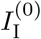 and period *T* the threshold condition (10) together with (9) determine *V*^(0)^(0). The periodicity condition (12) together with the jump condition (11) yield then the period *T* as a function of 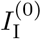. It is easier, however, to express 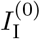 and *V*^(0)^(*t*) as a function of *T*. At 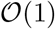 one then obtains

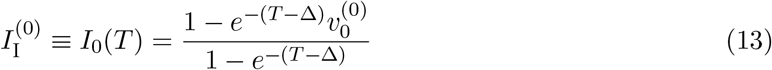

and (Fig.3a)

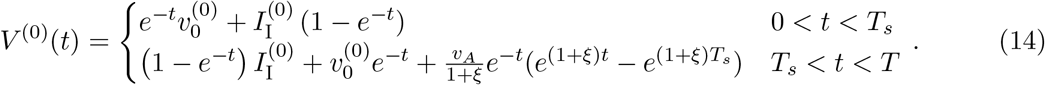

with

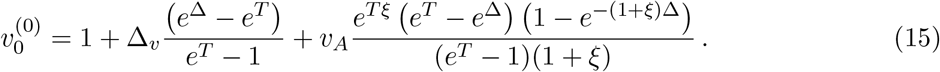

At 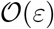 one obtains

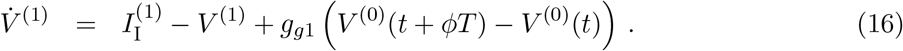

The threshold condition

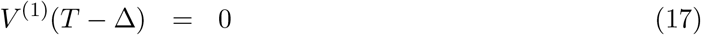

determines now the voltage *V*^(1)^(0), and the jump condition

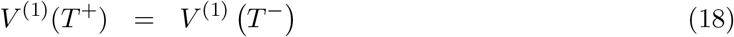

yields the correction 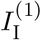 to the current that is needed to keep the period fixed. The dependence of 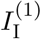 on the ISPD *ϕ* is of particular interest to us. The analytic expression for 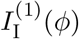 is too unwieldy to display and will be omitted here.

Considering a finite chain of unidirectionally coupled segments, *k* = 0,…, *N*, the expansion (7,8) can be viewed from a slightly different perspective. Since the leading neuron in segment *k* = 0 does not receive any gap junction input, its voltage is given by (14). Since all neurons have the same period, the period of the whole chain is therefore determined by the current into segment 0 via (13). The neuron in segment 1 receives synaptic input from the neuron in segment 0. Its voltage is therefore exactly given by *V*^(0)^(*t*) + *ϵV*^(1)^(*t*). The neuron in segment 2 receives synaptic input from neuron in segment 1 and its input includes therefore terms of 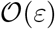 as well as 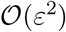. Analogously, neurons in subsequent segments receive inputs of multiple, including yet higher orders. Thus, a periodic wave initiated in the rostral segment *k* = 0 and traveling caudally on a semi-infinite chain of segments is given to 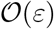 by

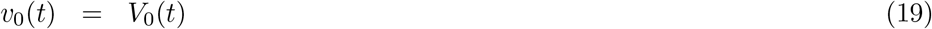

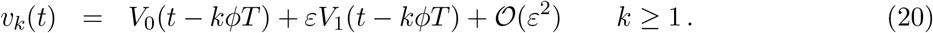

The reticulospinal input to neuron *k* = 0 is given by 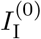, while that to the neurons with *k* ≥ 1 is given by 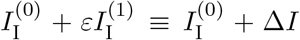. In the regime of interest the leading neuron at *k* = 0 is spiking without any additional input, while the follower neurons with *k* ≥ 1 also get the additional gap junction current from their rostral neighbor. Hence for positive ISPD *ϕ*, Δ*I* ≤ 0 and the overall reticulospinal input for follower neurons *k* ≥ 1 is lower than that of the leading neuron (i.e., *I*_*I*,0_ > *I_I,k_*). This interpretation of the perturbation expansion will guide us in the simulations in the remainder of this paper. Along these lines, Fig.(3a) shows the voltage traces of the V2a-I neurons in segment 0 and in segment 1 and also the contribution to the voltage from the gap junction current.

### 2.2 Direct Simulations

To assess the quality of the asymptotic approximation obtained in Section 2.1 we performed direct simulations of (1,3) for finite chains of length *N* = 8. The expansion turns out to be adequate even for quite sizable values of *g_g_* (Figure 4a). In the following we use the asymptotic approximation mostly as a guide for the numerical simulations. We therefore do not limit the values of *g_g_* to sufficiently small values that would yield quantitative agreement.

**Figure 4:**
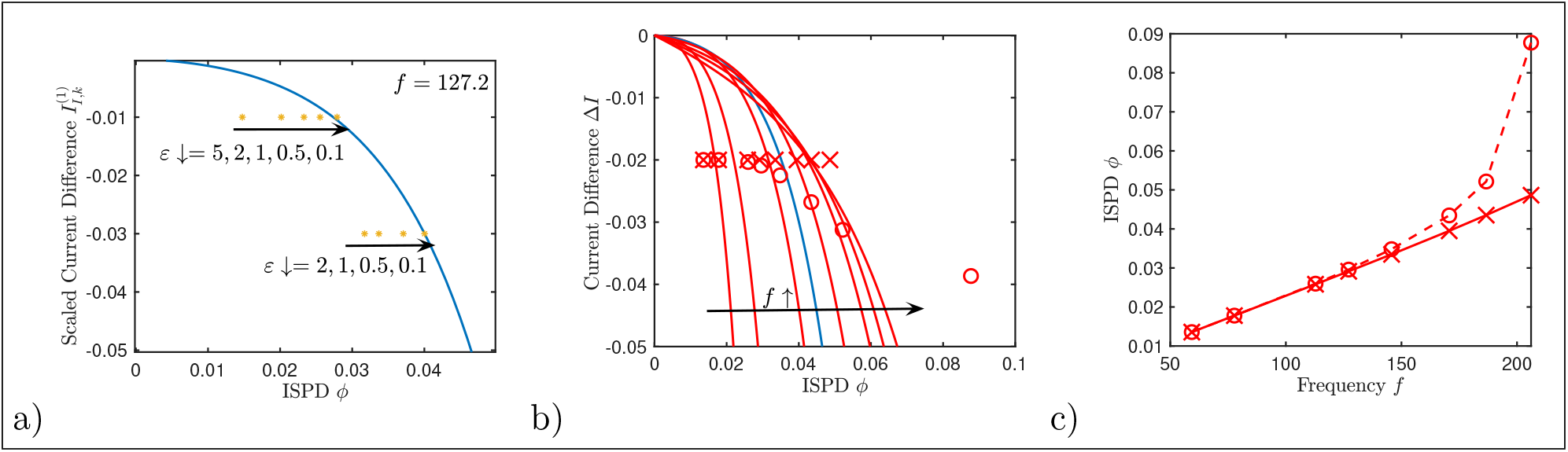
Intersegmental phase difference *ϕ*. (a) The numerical solution (yellow asterisks) converges to the asymptotic approximation (line) as *ε* → 0. *f* = 127.2, *g_g_* = *ϵ*, 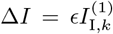. (b) Analytical results for Δ*I*(*ϕ*) for *f* = 59.3, 77.7,112.8,127.2,145.7,170.7,186.7, 206.7 (increasing to the right.) Scenario 1 (crosses): the base input for the whole chain, 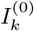, is increased with frequency, while Δ*I* is kept constant. Scenario 2 (circles): only the base input of the leader neuron is increased, implying that 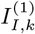 is reduced to keep *I_I_* constant. Gap junction coupling strength is fixed at *g_g_* = 1 for both scenarios. The ISPD varies less with frequency for scenario 1. (C) ISPD *ϕ* as a function of the frequency for scenario 1 (crosses) and scenario 2 (circles) with chain length *N* = 8.

Of particular interest is the dependence of the ISPD *ϕ* on the current difference Δ*I* and the frequency *f* of the wave. For sufficiently negative Δ*I* (Δ*I* ≲ −0.02 in Figure 4b) and not too high frequencies *f* the ISPD *ϕ* increases monotonically with *f* (Figure 4b). Since *ϕ* is defined as the spike-time difference relative to the period, this is the case when the time it takes a spike in neuron *k* - 1 to push the voltage in neuron *k* across the spiking threshold does not decrease sufficiently fast with increasing frequency. The gap junction current driving neuron *k* is proportional to the voltage difference between the two neurons. For fixed difference in the spike times of the two neurons that voltage difference increases with increasing slope of the voltage trace *v*(*t*). Since the slope increases with frequency, the gap-junction current increases, as well, decreasing the spike-time difference. For sufficiently large Δ*I* this leads to a decrease in *ϕ* for large frequencies (Δ*I* ≳ −0.02 in Figure 4b) and a non-monotonic *f*-dependence. In the following, we will be mostly concerned with the low-frequency regime where *ϕ* increases with *f*.

Ideally, the ISDP *ϕ* would not depend on the frequency of the wave at all in order to keep the undulation wavelength constant. This would require that Δ*I* be tuned quite precisely in parallel with any frequency changes. To avoid such a complex control task we first investigate the dependence of *ϕ* on the frequency of the wave in two simple scenarios (Fig.4b,c):

1. When changing the frequency, the synaptic reticulospinal input is adjusted uniformly for the whole chain, *k* ≥ 0, while the leader neuron receives additional, frequency-independent input. Thus, Δ*I* < 0 is kept fixed, while 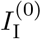 is adjusted to change the frequency.
2. When changing the frequency, the synaptic reticulospinal input *I*_I,*k*_ is changed only for the leader neuron *k* = 0. In this scenario Δ*I* is reduced as *I*_I,0_ is increased in order to keep *I*_I,*k*_ fixed for *k* ≥ 1 (cf.(13)).

For low frequencies both scenarios result in a very similar dependence of the ISDP on the frequency. For higher frequencies, however, the ISDP increases substantially faster with frequency in scenario 2, where only the input to the leading rostral neuron is increased (Fig.4c). From this perspective scenario 1 would be preferable.

However, in scenario 1 the total input to the follower neurons is increased with increasing frequency. For high frequencies this can bring them above threshold, *I*_I,*k*_ = *I*_I,0_ + Δ*I* > 1, allowing them to fire on their own without being triggered by their rostral neighbor. As a result, when swim motion is initiated from random initial conditions by a step in the input current, neurons in different segments are likely to fire out of order initially and it takes a number of oscillation periods for the firing of the chain to become ordered in a rostral to caudal manner. Thus, for *I*_I,*k*_ > 1 the initial swimming motion is expected to be disorganized. Indeed, the fraction of random initial conditions for which in simulations the correct firing sequence is established within 3 oscillation periods drops substantially when the input is increased above *I*_I,*k*_ = 1 (corresponding to the yellow dashed line in Fig.5). For weak gap-junction coupling the time to establish the correct firing sequence also becomes long.

**Figure 5:**
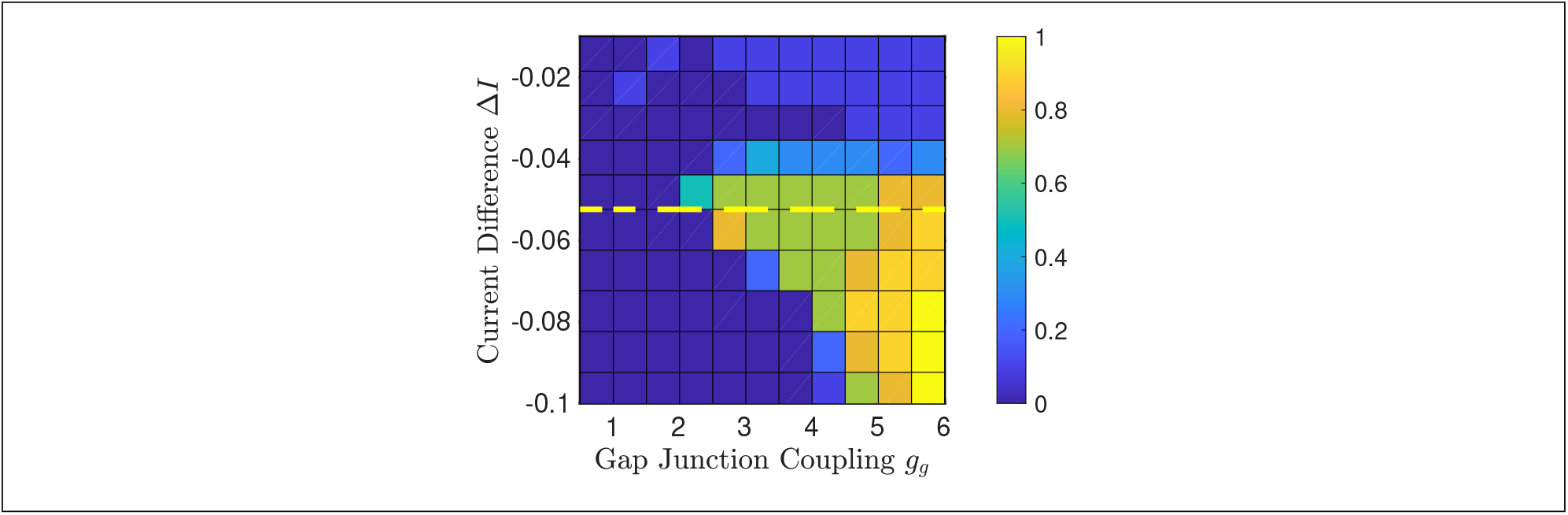
Establishing the correct spiking order. The color indicates the fraction of random initial conditions that reach the correctly sorted steady state within three oscillation cycles. The dashed line indicates *I*_I_ = 1. Parameters: *f* = 256, *N* = 8.

When Δ*I* is too negative for a given gap-junction coupling, no periodic wave can be established at all, i.e. the oscillations of the leader neuron may drive only a few of the more caudally located segments, but fail to trigger a wave that propagates all the way to the caudal end of the chain (solid lines in Fig.6). The propagation along longer chains requires Δ*I* to be less negative (dark vs light colored lines in Fig.6).

**Figure 6:**
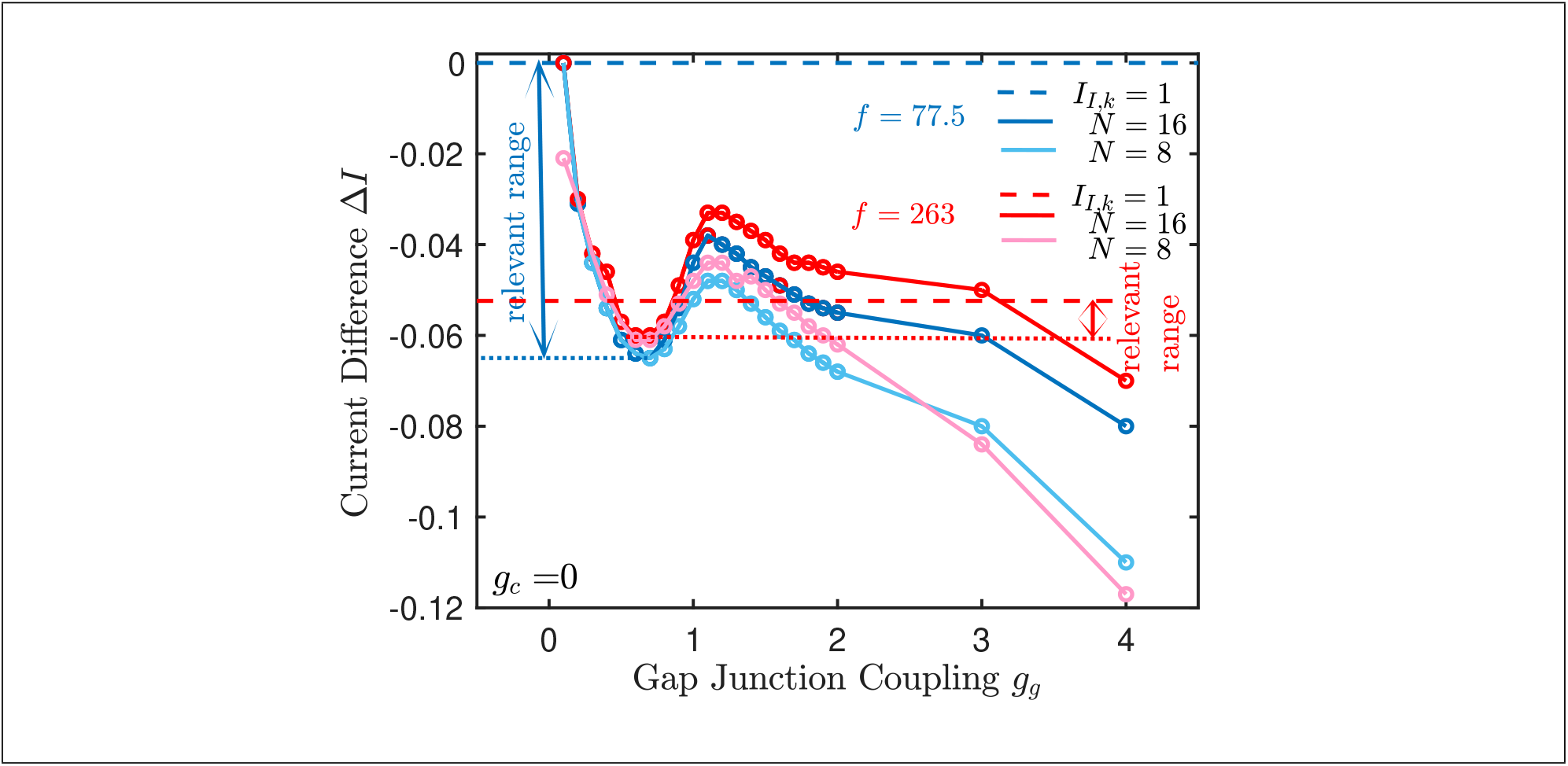
Range of relevant Δ*I* in scenario 1. Below the solid lines waves do not propagate throughout the whole chain of length *N* (*f* = 77.5 with *N* = 8 (light blue), *f* = 263 with *N* = 8 (pink) and *N* = 16 (red)). The dotted lines indicate the minimal value of Δ*I* if *g_g_* is restricted to *g_g_* ≲ 1. The dashed lines are a proxy for the upper bound beyond which the sorting becomes too slow (cf. Fig.5).

As mentioned before, we are interested in waves that satisfy two conditions: they propagate along the whole chain and their spiking order is quickly established from random initial condition. Fig.5 shows that *I*_I,*k*_ = 1 serves as a reasonable proxy for the maximal value of *I*_I,*k*_ for which the correct order is established sufficiently fast. The proxies for *f* = 77.5 and *f* = 263 are shown in Fig.6 as dashed lines. For *f* = 77.5 there is quite a range in Δ*I* for which both conditions are satisfied (blue double arrow). For *f* = 263, however, that range is restricted to large values of the gap-junction coupling, which are most likely unphysiological, and a very small range at low *g_g_* (red double arrow).

The range of physiologically sound and therefore relevant gap junction conductance can be estimated from the coupling coefficients measured between V2a-I neurons. The coupling coefficients between V2a-I neurons that are 2 to 4 segements apart were recorded to be in the range 0.005-0.02 (Menelaou and McLean, 2019). This suggests a coupling coefficient up to 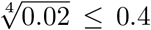. In our model the coupling coefficient for directly connected neurons can be derived as 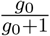. The measurements thus suggest that the maximum gap junction conductance *g*_0_ is less than 2/3 and we will accordingly focus on gap junction conductance values less than 1. Thus, in this minimal model the physiologically sound range of gap junction conductance allows reasonable swimming motion (short transient period and reliable spike propagation) at the higher swimming speed only in a very narrow range of Δ*I* and *g_g_*. In the following section, we therefore investigate how the feedforward synaptic connections mediated by V2a-II neurons can expand the valid range of Δ*I* and discuss how to keep the ISPD relatively constant over a range of frequency.

## 3 Interleaved Chains of V2a-I and V2a-II Neurons

The two types of V2a interneurons exhibit quite distinct morphology and electrophysiology. The resulting functional differences are not yet fully understood. Here, we explore the possible role of V2a-II neurons in the rhythmic control of the spinal cord network and add V2a-II neurons to the model described above. It has been observed that the V2a-II neurons tend to spike earlier on average than the V2a-I neurons (Menelaou and McLean, 2019). In our modeling this is in part due to their electrophysiological differences and in part to their different connectivities.

V2a-II neurons show significant spike-frequency adaptation (Menelaou and McLean, 2019). To capture this behavior we extend the neuron model (1) to include an adaptation current *w*,

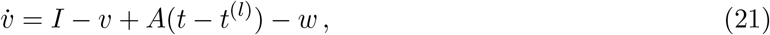

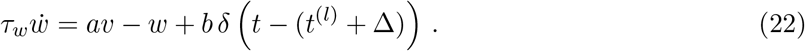

The Dirac *δ*-function incorporates a reset of *w* at *t* = *t^(l)^* +Δ, i.e. at the end of the action potential. Due to the adaptation, the initial spike frequency is twice as a large as the tonic frequency and substantially contributes to the early spiking of the V2a-II neurons compared to the V2a-I neurons (Fig.7).

**Figure 7:**
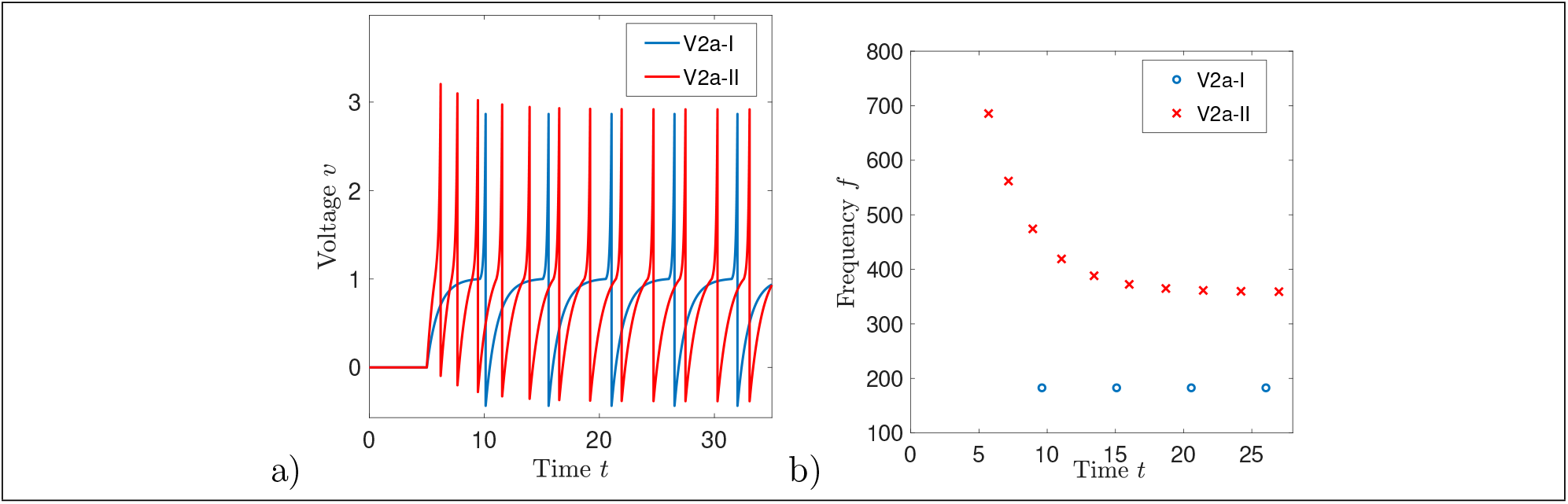
Spike-frequency adaptation of V2a-II, but not of V2a-I neurons. (a) Voltage traces for V2a-I (blue, cf. (1)) and for V2a-II (red, cf. (21,22)). b) Instantaneous frequency defined via the inter-spike intervals for step-current input for V2a-I and V2a-II neuron.

V2a-I and V2a-II neurons project caudally to the respectively other type of neurons. Unlike the V2a-I to V2a-I connections, the connections between V2a-I and V2a-II are, however, predominantly *via* chemical (glutamergic) synapses (Menelaou and McLean, 2019). For simplicity, we will assume that these chemical synaptic projections only reach to the neighboring caudal segment. Thus, the V2a-I and V2a-II neurons form an interleaved network of the kind shown in Figure 1. Note that in our model the V2a-II neuron segment 0 receives input from the V2a-I neuron in the same segment instead of the previous (non-existent) segment. The V2a-II neurons also receive reticulospinal input similar to V2a-I neurons, which we denote by *I*_II_. In this network the V2a-I neurons receive input from both V2a-I and V2a-II neurons of their rostral neighbor segment. The input from the V2a-I is through a gap junction, while that from the V2a-II is via a chemical synapse. Moreover, the V2a-II neuron receives its input from the V2a-I neuron in its rostral neighbor segment via a chemical synapse. Therefore, the relative timing of the inputs to the caudal V2a-I neuron *via* the gap junction and *via* the chemical synapse can be controlled by changing, e.g., the bias current *I*_II_ of the V2a-II neurons.

To model the chemical synapse we add for each spike at time 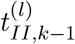 of the presynaptic V2a-II neuron a conductance-based synaptic current with reversal potential *E_syn_* to (1),

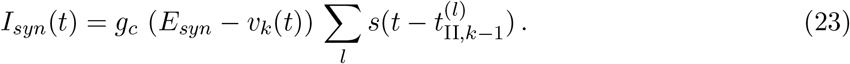

The current is assumed to have a fixed waveform given by the difference between two exponential functions,

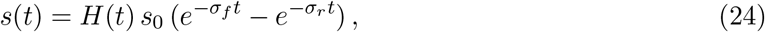

with *H*(*t*) denoting the Heaviside step function. With the normalization factor *s*_0_ = *σ_f_ σ_r_*/(*σ_f_* – *σ_r_*) the maximal conductance is *g_c_*.

Since the chemical synapse is excitatory (*E_syn_* > *V_T_* = 1), it provides additional depolarizing input to the V2a-I neurons (Fig.8). This input has a different effect than increasing the tonic base input *I_I_*, since it is active only over a short time window, the timing of which is controlled by the upstream V2a-II neuron. Despite the additional excitation, this allows the V2a-I neurons to stay below threshold until their rostral neighbor has spiked. As a result, the range of Δ*I* over which the wave propagates reliably across all *N* segments is increased without significantly affecting the maximal value of Δ*I* that allows sufficiently fast ordering (compare Figs.9,10 with Figs.5,6). Thus, the relevant range of Δ*I* is significantly increased by the chemical synapse. In fact, while without V2a-II input no values of Δ*I* (except for a very narrow range) are relevant for gap-junction strengths less than *g_g_* = 1 (for *N* = 8) and *g_g_* = 3 (for *N* = 16), with chemical synapses there is a substantial relevant range of Δ*I* for all values of gap-junction strength we are interested in.

**Figure 8:**
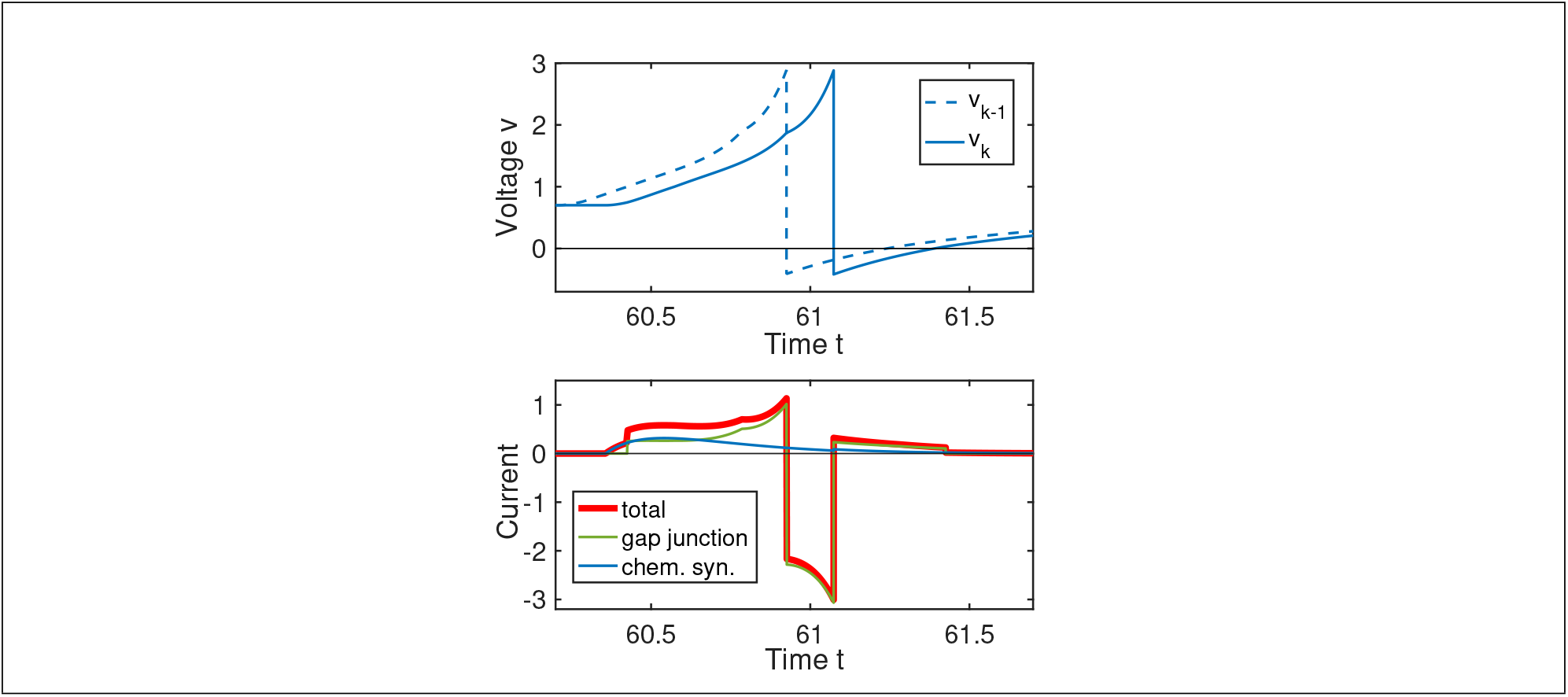
Inputs to the V2a-I neurons via chemical synapses and gap junction. Top: voltages *v*_*k*-1_ and *v_k_* of V2a-I neurons *k* (solid) and *k* – 1 (dashed). Bottom: total synaptic input current (red, thick) arising from a chemical synapse (blue), which is driven by a spike in a V2a-II neuron that is triggered by the spike of V2a-I neuron *k* – 1, and a gap junction (green) between the V2a-I neurons *k* – 1 and *k*.

**Figure 9:**
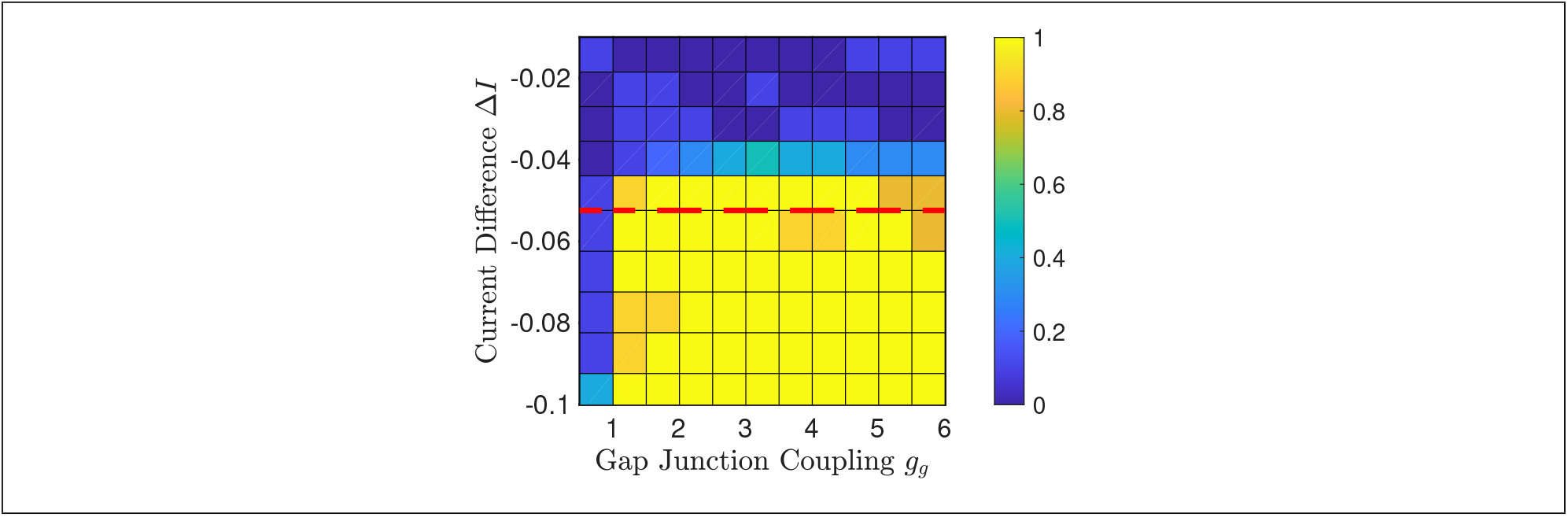
Sorting with V2a-I neurons coupled to V2a-II neurons *via* chemical synapse. (cf. Fig.5). The color indicates the fraction of random initial conditions that reach a correctly sorted steady state within three cycles. *f* = 256, *N* = 8, *g_c_* = 0.01 and *I_II_* = 1.1. The red dashed line indicates *I*_I,*k*_ = 1.

**Figure 10:**
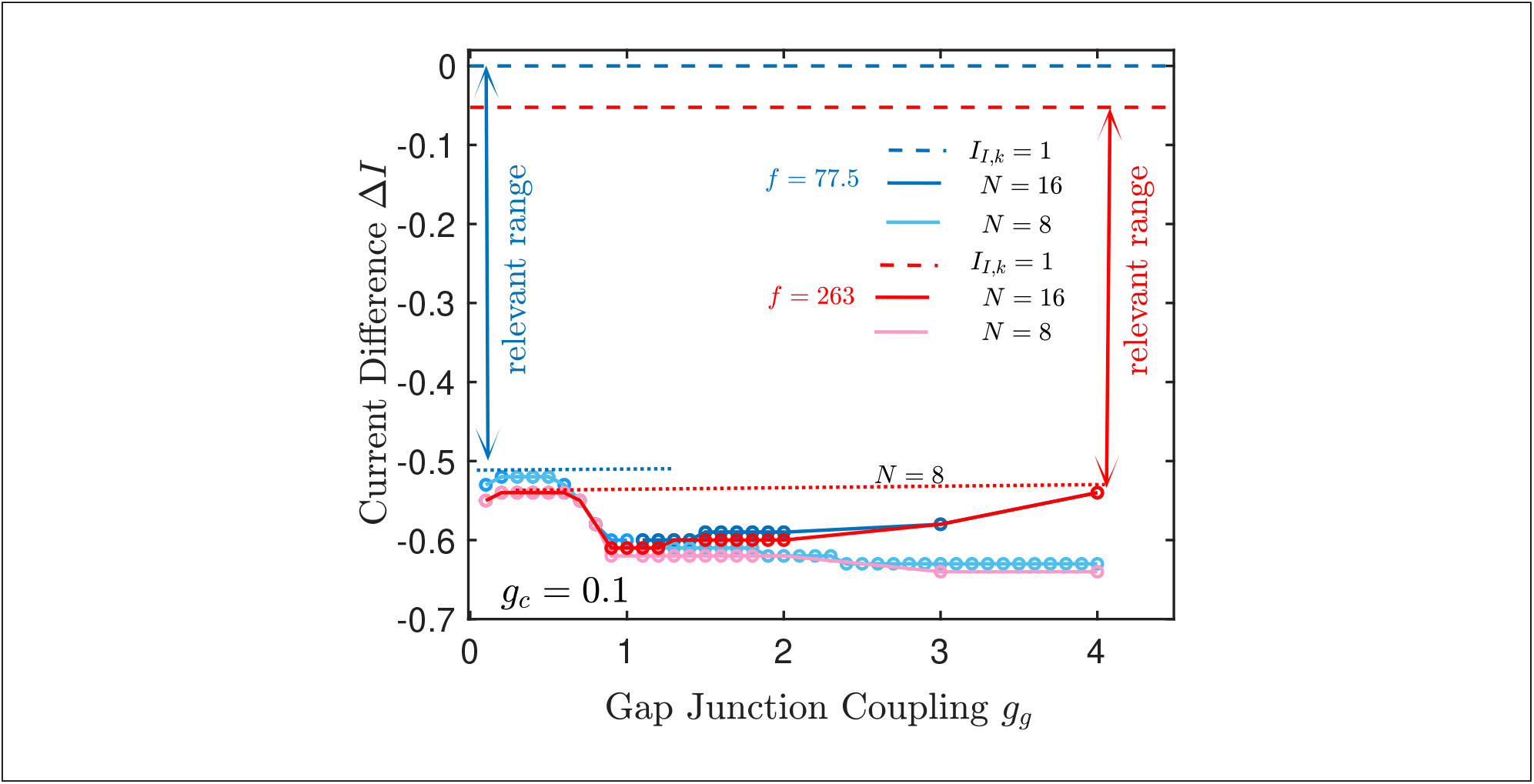
Range of relevant values of 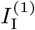 with V2a-II neurons. As in Fig.6, solid lines denote the lower limit for Δ*I* for waves to propagate throughout the system (*f* = 77.5 with chain length *N* = 8 (light blue) and *N* =16 (blue), and *f* = 263 with *N* = 8 (pink) and and *N* = 16 (red)). The dotted lines indicate the lower limits when *g_g_* is limited to *g_g_* ≲ 1. The dashed lines denote the proxies for the upper limits for fast ordering. The double arrows indicate the relevant range of Δ*I* for *f* = 77.5 and *f* = 263, respectively. The base input level of V2a-II neuron was fixed at *I_II_* = 1.4.

### 3.1 Control of *ϕ* by V2a-II Neurons

To gain insight into the impact of the chemical synapses on the ISPD *ϕ* between the V2a-I neurons we include them in our asymptotic calculations and consider their synaptic input from the rostral V2a-II neuron to be of the same order as the gap junction current from the rostral V2a-I neuron. At 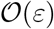 the equation in the asymptotic expansion can then be written as (cf. (23))

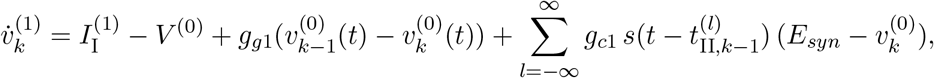

where *E_syn_* is the reversal potential of the chemical synapse. Note that the conductance depends on the time 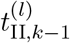 at which the V2a-II neuron in segment *k* – 1 reaches the spike threshold, which in turn depends on the spike time of the preceding V2a-I neuron in segment *k* – 2. We express the spike time of the rostral V2a-II neuron relative to the spike time of the preceding ((*k* – 1)^*th*^ segment) V2a-I neuron and denote their time difference as *τ*_Δ_. Focusing on periodic solutions with period *T* that are phase-shifted by *ϕ* between consecutive segments, we obtain then

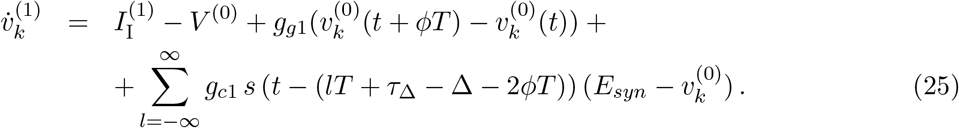

As in the case without chemical synapses, these equations can be solved analytically, but the resulting expressions are too involved to display explicitly. Again we use the analytical results as guidance for the direct simulations without aiming for quantitative agreement.

While the frequency range over which waves propagate robustly along the chain of V2a neurons is increased with the addition of V2a-II neurons (cf. Figs.6,10), the ISPD *ϕ* still increases significantly with increasing frequency (Fig.11a,b,c). Thus, for fixed Δ*I* the undulating swimming pattern of the animal still changes with swimming speed (cf. Fig.4b). However, if we allow Δ*I* to become less negative with increasing frequency, the range in *ϕ* can be reduced. The minimal and maximal values of *ϕ* are then determined by the range of relevant Δ*I* values (cf. Fig.10), which is marked by the solid portion of the thick lines in Fig.11a,b,c. Thus, the minimal variation in *ϕ* is given by the difference between the maximal *ϕ* at the minimal frequency and the minimal *ϕ* at the highest frequency. Without chemical synapses the ISPD varies then at least from 0.029 to 0.041 across the frequenc range 77.5 ≤ *f* ≤ 204 (blue lines for perturbation results and diamonds for numerical results in Fig.11a).

**Figure 11:**
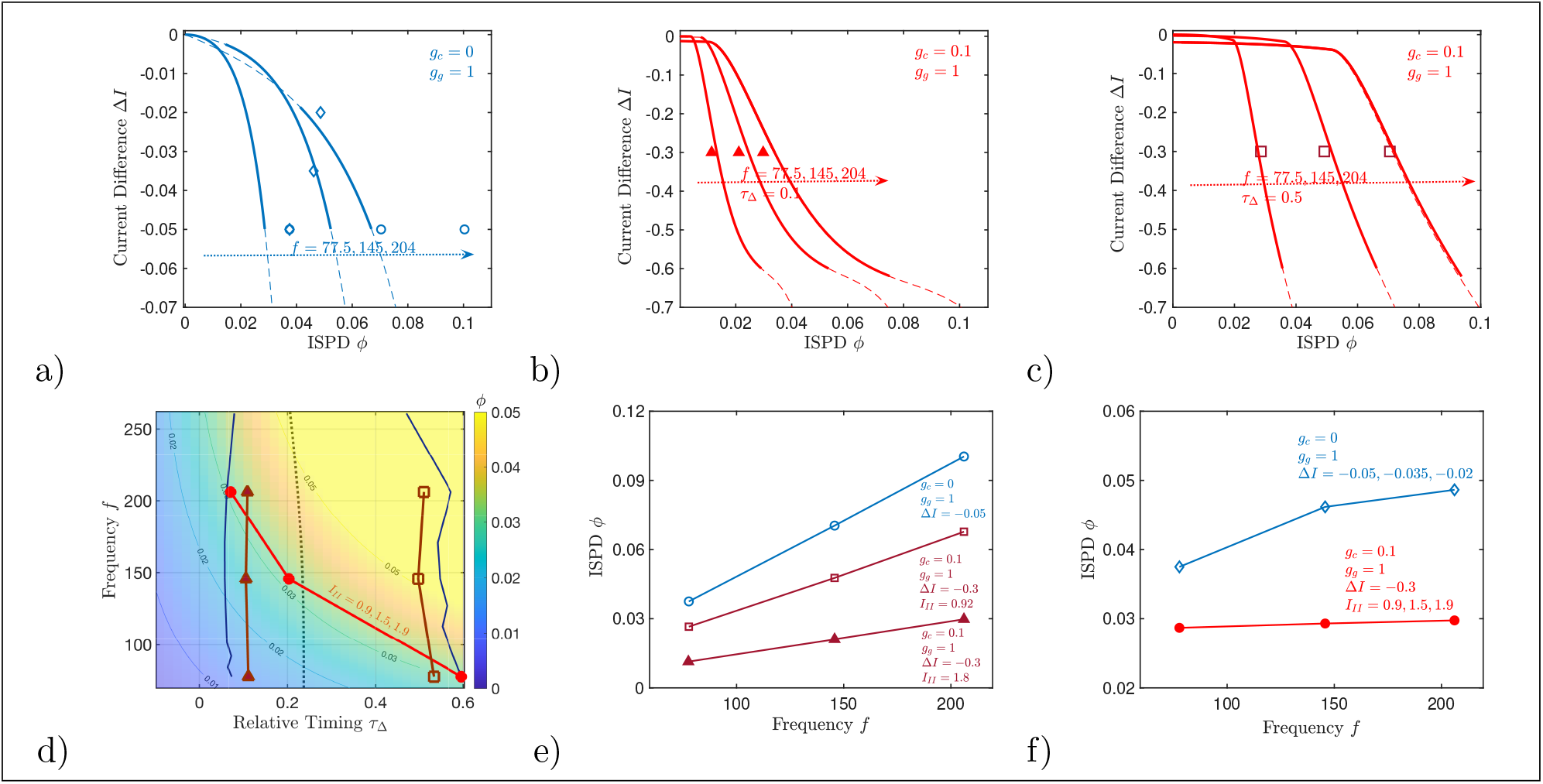
Frequency-dependence of the ISPD *ϕ*(*f*). (a-c) Asymptotic approximation Δ*I*(*ϕ*) for different frequencies (lines). The relevant range of Δ*I* is indicated by the solid lines: above that range the propagation of spikes along the chain is not reliable and below that range sorting is too slow (cf. Figs.6,10). Diamonds, circles, triangles, and squares refer to numerical simulations, which are also shown in (d,e,f). (a) only gap junctions between V2a-I neurons (*N* = 8), (b,c) Lines: with additional chemical synapses between the V2a-II and the V2a-I neurons (*N* = 16) for *τ*_Δ_ = 0.1 and *τ*_Δ_ = 0.5, respectively. The symbols denote the same simulations as in panel (d) with *I*_II_ = 0.9 and *I*_II_ = 0.45 yielding *τ*_Δ_ ≈ 0.1 and *τ*_Δ_ ≈ 0.5, respectively. (d) ISPD *ϕ* as a function of the frequency *f* and the delay *τ*_Δ_ for Δ*I* = −0.3 and *g_c_* = 0.1. To obtain values for *τ*_Δ_ to the left of the left blue line *I*_II_ would have to be chosen so large that the V2a-II spike irregularly and more than once per cycle, while to the right of the right blue line they do not spike at all. The dark red lines correspond to simulations with fixed *I*_II_ = 1.8 (triangles) and *I*_II_ = 0.92 (squares). The dashed black line marks *ϕT* = *τ*_Δ_. Δ*I* = −0.3. (e) ISPD *ϕ*(*f*) from numerical simulations for fixed Δ*I* without (blue) and with (red) chemical synapses (same simulations as in panels (a,d)). (f) Without V2a-II, the variation of *ϕ*(*f*) can be minimized by tuning Δ*I* (blue diamonds). With V2a-II, tuning the tonic input *I*_II_ allows *ϕ* to be kept identically constant (red circles).

While the minimal *ϕ* at the maximal frequency is determined by the upper limit of Δ*I*, which is not affected by the V2a-II neurons, the input from V2a-II neurons substantially reduces the lower limit of Δ*I*. As a result, the *ϕ*-values associated with the relevant range of Δ*I* can overlap significantly for different frequencies. For instance, for *τ*_Δ_ = 0.1 the ISPD can be held constant over the frequency range 77.5 ≤ *f* ≤ 204 for any value of *ϕ* within the range 0.01 ≤ *ϕ* ≤ 0.03 by tuning Δ*I* (Fig.11b). Such a range in the ISPD exists also for larger *τ*_Δ_ (Fig.11c).

The ISPD *ϕ* depends significantly on *τ*_Δ_, as shown for fixed Δ*I* in Fig.11d. For fixed *I*_II_, *τ*_Δ_ changes slightly with frequency, contributing minimally to the frequency dependence of *ϕ* (dark red symbols in Fig.11d,e). Note that with decreasing *τ*_Δ_ the frequency-dependence of *ϕ* becomes weaker. Since increasing *I*_II_ leads to earlier V2a-II firing, i.e. smaller *τ*_Δ_, it allows to render *ϕ* independent of frequency by tuning *τ*_Δ_ via *I*_II_ (bright red open circles in Fig.11d,f). However, tuning the base input level *I*_II_ precisely enough when varying the frequency may not be physiologically realistic. Moreover, the range of *τ*_Δ_ is limited by the range of *I_II_* that allows reliable spiking of V2a-II neurons for given frequency *f* (blue lines in Fig.11d). We therefore investigate in the next section the simpler strategy of having the V2a-II neurons spike only for higher frequencies.

The spikes of the V2a-I and V2a-II neuron in a given segment are triggered by the same V2a-I neuron in the upstream neighboring segment. Relative to the spikes of that neuron their spike times are delayed by *ϕT* and *τ*_Δ_, respectively; they spike simultaneously along the dashed line *ϕT* = *τ*_Δ_ in Fig.11d. When increasing the frequency, *τ*_Δ_ needs to be decreased in order to keep *ϕ* constant (bright red open circles in Fig.11d). Thus, for larger frequencies *τ*_Δ_ needs to be negative, i.e. the V2a-II neuron needs to fire before the V2a-I neuron in the same segment. Since the chemical synapse driving the V2a-II neuron introduces a slight delay compared to the gap-junction driving the V2a-I neuron, this might be unexpected. However, in experiments the V2a-II neurons are indeed observed to fire before the corresponding V2a-I neurons (Menelaou and McLean, 2019).

### 3.2 Spike-Frequency Adaptation of V2a-II Neurons and *ϕ* Control

For low values of *τ*_Δ_ the additional input from the V2a-II neurons reduces the ISDP *ϕ* of the wave. Since *ϕ* increases with frequency it is expected that the overall variation of *ϕ* with varying frequency can be reduced by having the V2a-II neurons being active only in the upper frequency range. An interesting question is therefore, whether the activity of the V2a-II neurons can be restricted to higher wave frequencies without any tuning of the base input level of V2a-II neurons with frequency.

In the scenarios considered here both types of V2a neurons spike only once per cycle. The activation of V2a-II neurons for higher frequencies can therefore not result from the integration of multiple fast inputs into the V2a-II. The neuron can therefore not be a simple integrator like (1). This activation can, however, be the consequence of appropriate timing of the inputs if the V2a-II neurons are resonators (Izhikevich, 2003). In fact, V2a-II neurons exhibit significant spike-frequency adaptation (Menelaou and McLean, 2019), which we have incorporated by extending the minimal neuron model (1) to include an adaptation current (21,22). For the synaptic strength used so far (*g_c,II_* = 0.1) the V2a-II neurons spike over the whole range of frequencies investigated. We consider now weaker synaptic strengths (*g_c,II_* = 0.055) for which the spiking depends on the timing of the inputs and therefore on the frequency of the wave.

A key feature arising from the adaptation current is the overshoot in the voltage that occurs during the recovery phase after a spike (arrow near *t* = 30.5 in Fig.12a). Synaptic input from the V2a-I neuron arriving during that overshoot can drive a spike in the V2a-II neuron (blue triangles in (Fig.12b), even if it is too weak to elicit a spike at a later time (red triangles). Thus, in this range of the synaptic input strength V2a-II neurons will spike and reduce the ISDP *ϕ* only in for larger swimming frequencies for which the synaptic inputs arise during the overshoot. The overshoot does not arise in the relevant time window if the previous input to the neuron only depolarized it without triggering a spike. A synaptic input that was strong enough to trigger a spike during the overshoot may then not be sufficient to activate a spike, even if it arises early (blue triangle in Fig.12c). Thus, this model of the V2a-II neuron exhibits significant hysteresis: once a spike is triggered, inputs with sufficiently high frequency maintain the spiking, but they will not initiate a transition to spiking if the frequency of the inputs is slowly increased. However, spiking will cease, even when the input frequency is decreased slowly.

**Figure 12:**
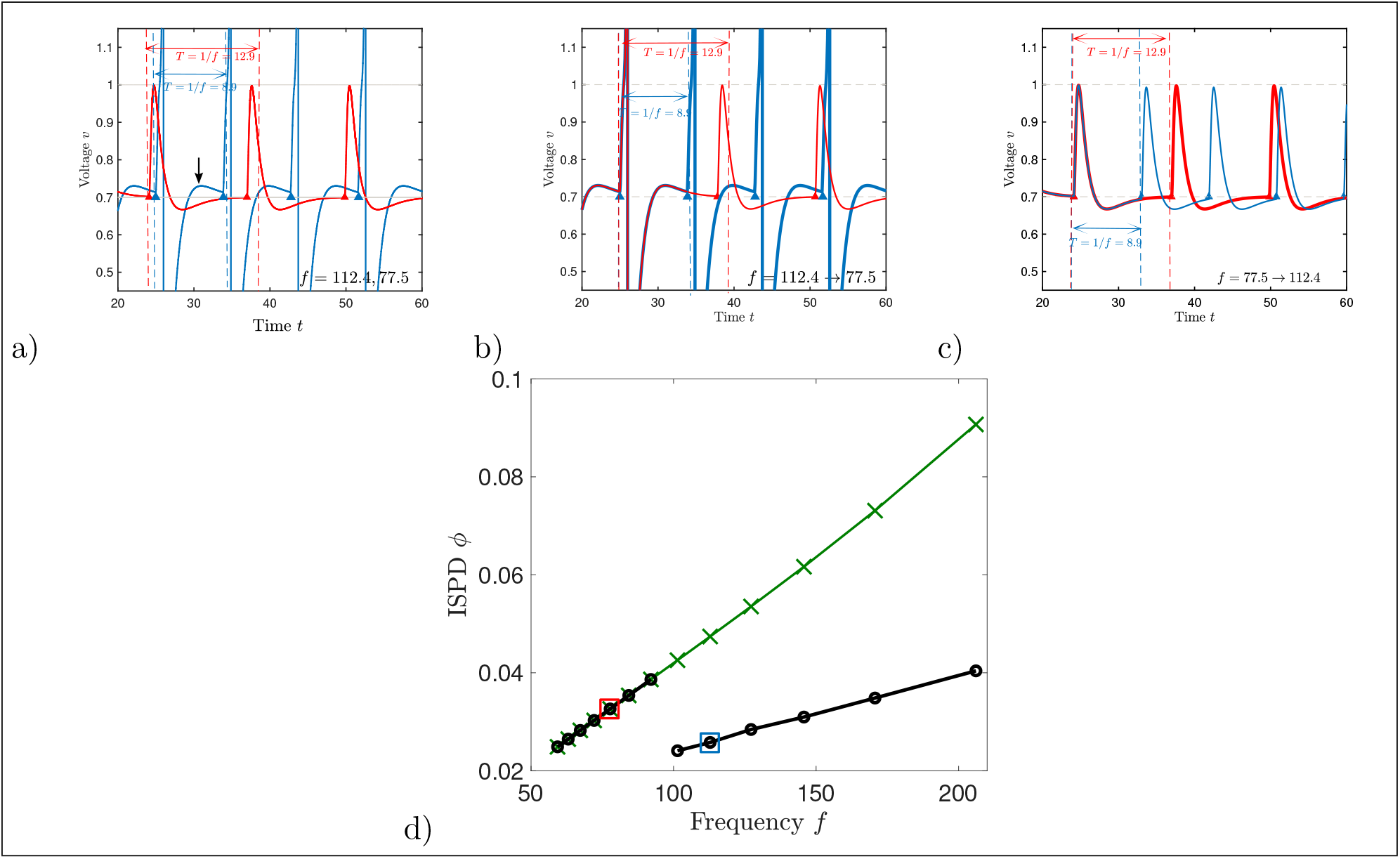
Adaptation of V2a-II neuron and *ϕ* control. Numerical solutions for V2a-II neurons with varying input frequency. *g_c,II_* = 0.055, Δ*I* = −0.04. a) Bistabiliy between no-spiking for low-frequency periodic input and spiking for high-frequency input. Arrow marks the peak of the overshoot. b) When switching the input from high to low frequency the V2a-II stops firing (red line). c) Switching from low to high frequency does not trigger spiking in the V2a-II (blue line). d) The variation of the ISPD *ϕ* with frequency is considerably reduced by chemical synaptic input from a V2a-II that responds only to high-frequency input.

Due to its frequency-dependent spiking the adaptive V2a-II neurons can reduce the variation of the ISDP *ϕ* with frequency. While without the V2a-II neurons *ϕ* increases substantially with increasing frequency (green crosses in Fig.12d), with the V2a-II neurons included the variation in *ϕ* is much smaller (black circles in Fig.12d), since *ϕ* is shifted down significantly once the V2a-II neurons are driven to spiking by the rostral V2a-I neuron spikes. This occurs above a certain frequency (*f_th_* ≈ 91.2 in Fig.12d). In these simulations initially none of the neurons receive any reticulospinal input, which implies low voltage and adaptation variable. Then the reticulo-spinal inputs are turned on for both V2a-I and V2a-II. Only the tonic input to the V2a-I neurons is varied to change the frequency of the wave; the input to the V2a-II neurons is the same for all frequencies. If the frequency of the wave was increased slowly from low values the V2a-II neurons would not start spiking once the frequency *f_th_* is reached due to the strong hysteresis. Consequently, the ISPD would not be reduced (green crosses in Fig.12d). However, appropriate spiking could be assured by including a brief boost in the base input *I*_II_ whenever the frequency is changed. This would trigger an initial spike of the V2a-II neuron that would be sufficient to initiate a transition to the spiking branch with its lower *ϕ* for high frequencies (black circles in Fig.12d), but not for low frequencies (Fig.12a,b).

Thus, the spike-frequency adaptation of the V2a-II neurons provides a mechanism that selectively provides additional current to V2a-I neurons during fast swimming. The ISPD still increases with frequency in both dynamic regimes, but the downshift of the ISPD when the V2a-II neuron becomes active reduces the overall variation of *ϕ* substantially. Based on this model one may surmise that having a variety of V2a-II neuron populations that switch on at different frequencies might decrease the range of ISPD further, as each activated V2a-II neuron will provide additional input that can reduce the time it takes the V2a-I neuron to become activated.

## 4 Discussion

This paper has been motivated by a recently proposed new model for the organization of the spinal cord of zebrafish larvae (Menelaou and McLean, 2019). Classically, two types of connectivity models for such motor circuits have been proposed. In the hierarchical, multi-layer model two separate interneuron populations make the higher-order connections receiving control signals from the brain and the last-order connections to the motorneurons, respectively (McCrea and Rybak, 2008; Rybak et al., 2015). In the single-layer model there is only a single interneuron population that maintains both types of connections (Kozlov et al., 2009). The experimental results of Menelaou and McLean (2019) suggest a hybrid combination of these two models, since it identified two separate, but interacting populations of excitatory interneurons (V2a-I,II) that each make last- and higher-order connections, albeit to different degrees. In particular, V2a-I neurons make weaker last-order connections to the motor pool than the V2a-II neurons. The roles that these different populations play in generating swimming motion have not been clarified yet.

Our minimal model of such a hybrid axial circuit suggests that the synergetic interaction of the two types of interneurons may serve two objectives. When the V2a-I neurons in different segments are coupled only via the electrical gap junctions, the rostral, leading V2a-I neuron evokes spikes only in a limited number of more caudally located neurons unless the gap-junction coupling is unphysiologically strong. Moreover, for more moderate coupling driving an undulation wave across ~ 30 segments requires tonic reticulospinal input to the caudal V2a-I neurons that is so large that the caudally located neurons can almost spike on their own without the input from their rostral neighbors. As a consequence, when starting from random initial conditions it takes quite a long time to establish the spiking order that corresponds to a proper undulation wave. Appropriately timed synaptic input from the V2a-II neurons to the V2a-I neurons overcomes this limitation. It allows propagating-wave solutions in long chains even when the tonic input to the V2a-I neurons is weak enough not to interfere with the rapid establishment of the correct spiking order along the segments.

In addition, the synaptic input from the V2a-II neurons to the V2a-I neurons modifies the ISPD depending on the timing of synaptic input relative to the gap-junction input the V2a-I neurons receive from their neighboring V2a-I neurons. This timing depends on the tonic input to the V2a-II neurons. The ISDP can therefore be kept constant by a suitable tuning of the tonic input to the V2a-II neurons. Less precise, but more robust and requiring very little tuning is a binary control scheme in which the V2a-II neurons are only spiking in the upper range of swimming frequencies. This aspect emerges naturally through the spike-frequency adaptation that the V2a-II neurons exhibit (Menelaou and McLean, 2019): there is a limited time window after each spike in which the V2a-II neurons are more excitable than at a later time. If the wave frequency is too low, the rostral inputs come too late and do not elicit any V2a-II spikes.

A characteristic feature of our model is the cooperation of chemical synapses and electrical gap junctions, which are assumed to be unidirectional based on the experimental results (Menelaou and McLean, 2019). Weak, but bidirectional coupling in long chains of coupled oscillators has been discussed comprehensively within the general phase-description framework that is possible for weak coupling (Kopell and Ermentrout, 1986; Kopell et al., 1991). There the range of existence and stability of phase-locked solutions was established and it was shown that typically the coupling in one direction dominates, which then determines the phase shift between the individual oscillators. In particular, it was shown that the phase shift is frequency-independent, if the coupling functions, which give the change in frequency of an oscillator as a function of the phase (rather than timing) difference between adjacent oscillators, do not depend on the oscillation frequency (Kopell and Ermentrout, 1986). This special behavior was explicitly obtained in a connectionist rate-type model for the lamprey spinal cord (Williams, 1992). As our results confirm, in general the frequency independence is not to be expected without tuning and an observation of a vanishing or weak dependence of the ISPD on the frequency points therefore to some special properties of the circuit.

To capture the neuronal interactions via gap junctions it is essential that the temporal trace of the action potential is included sufficiently well in the treatment. In order to make analytical progress we used a model put forward by Chow and Kopell (2000) in their detailed analytical and numerical investigation of bi-directional gap-junction coupling. There the spiking currents are modeled by inhomogeneous, explicitly time-dependent terms, which allows the equations to remain linear. In contrast to our analysis, their work focused on all-to-all coupling and identified, in particular, the role of the spike shape on the formation of synchronous and anti-synchronous states as well as splay-states, which are akin to unforced traveling waves. This approach ignores the change in the shape of the action potentials of the interacting neurons that results from the interaction. This becomes, however, significant only for stronger coupling when the current induced by the coupling becomes of the order of the spiking currents themselves. An exact treatment of the gap junction coupling that includes changes in the wave forms is possible in a population description of heteregeneous quadratic-integrate-fire neurons (Pietras et al., 2019).

Multi-layer and single-layer models differ in characteristic ways in their response to the deletion or inhibitory suppression of specific neurons. In single-layer models the inhibition of the last-order neurons, which give output to the motor pool, suppresses the whole rhythmic activity since these neurons are also providing the timing information. Brief such suppressions reset the rhythm and are therefore called resetting deletions. In contrast, in multi-layer models the inhibition of the last-order neurons does not affect the higher-order neurons and has no effect on the rhythm (‘non-resetting deletion’). In the hybrid model, the suppression of V2a-II neurons does not suppress or reset the rhythm. However, it suppresses the input to the V2a-I neurons via the chemical synapse and therefore increases the ISPD between the segment in which the suppressed neuron is located and its caudal neighbor. Thus, relative to the most rostral segment the phases of all segments caudal to the suppressed neuron are shifted by the same amount. If V2a-II neurons are suppressed in multiple segments the phase shift of the more caudally located segments is proportional to the number of suppressed neurons. Measurements of the phase shift resulting from suppressing neuronal spikes should allow an experimental assessment of the role of the chemical synapses between V2a-I and V2a-II neurons in the timing control during swimming and would provide a test of the model. In multi-layer models higher-order neurons are typically associated with timing control, while last-order neurons are assumed to play a key role in controlling the amplitude of the motion. Thus, increasing tonic reticulospinal input to the last-order neurons can increase the number of spikes elicited by each timing pulse emitted by the higher-order neurons, thus increasing the amplitude of the motion. The increased tonic input will also affect the timing of the spikes in the last-order neurons, but if they all receive the same reticulospinal input that phase shift would be the same for all segments and the ISPD would be unaffected. Thus, the wavelength of the undulation would be independent of the amplitude. This independence is not likely to persist in the hybrid model. Even though it features two different excitatory populations, they interact with each other via chemical synapses. In the hybrid model discussed here the same global change of the reticulospinal input leads to a similar change in the spike timing of the V2a-II neurons and with it of the input to the V2a-I neuron *via* the chemical synapse. Since that input modifies the ISPD between the V2a-I neurons, a global change in the reticulospinal input induces a change in all ISPDs. Thus, in contrast to the wavelength of the multi-layer model, the wavelength of the hybrid model investigated here varies with the amplitude of the motion. Based on the regime considered here, in which the ISPD decreases with increasing tonic input to the V2a-II neurons, the model predicts that at fixed swimming frequency stronger swimming motions would be associated with longer wavelengths of the undulation.

The experimental observations suggest that in zebrafish the V2a-I neurons resemble more closely first-order neurons, while the V2a-II neurons are more akin to last-order neurons. What distinguishes this network from a hierarchical model are the connections from the V2a-II neurons back to the V2a-I neurons. In our computational model these connections play a key role in enabling the network to support reliably and properly ordered propagating waves that can be established quickly from random initial conditions with ISPDs that do not depend strongly on the swimming frequency. While these requirements are likely to be very important for undulatory swimming motion, they are less relevant for legged locomotion. This could explain why the zebrafish spine has this hybrid structure, while the spinal networks of four-legged animals have a hierarchical structure (Rybak et al., 2015).

In the minimal hybrid model discussed here the V2a-II neurons fire only a single action potential in each cycle. This would only allow rudimentary amplitude control. For low reticulospinal input the V2a-II neurons do not spike. Nevertheless, the motor pool is driven in the hybrid model, but solely by the (somewhat weaker) inputs from the V2a-I neurons (Menelaou and McLean, 2019). For larger reticulospinal input the V2a-II neurons spike and substantially increase the drive to the motor pool. At the same time, the spike-driven synaptic input to the V2a-I neurons affects the ISPD. One could envision a more refined model in which the V2a-II neurons fire in bursts with a duration that increases with reticulospinal input. How such a change in amplitude would affect the ISPD would depend on the timing of these spikes. At this point the encoding of the amplitude in the V2a-II activity pattern is not sufficiently well known to warrant a detailed model capturing this aspect.

In this study we have only considered simple periodic swimming activity. An important component of that motion is its left-right alternation. We expect that this can be captured by a straightforward extension, as it has been done in many previous studies (e.g. Kozlov et al., 2009), in which corresponding populations on the left and right side inhibit each other via interneurons like the V0d neurons (Menelaou and McLean, 2019). The role of the various neuronal populations in other, more complex motions like turning, orienting, or self-righting remains an interesting open question.

## Acknowledgments

We gratefully acknowledge discussions with D.L. McLean and funding by NSF (DMS-1547394) and NIH (DC015137).

